# Assessment of the Role of Telocyte Zbtb16 in Lymphatic Drainage using Efhd1-CreERT2 x Zbtb16^flox/flox^ Mice

**DOI:** 10.1101/2025.11.28.691161

**Authors:** Yue Peng, Edward M. Schwarz

## Abstract

Lymphatic function is critical for maintaining fluid homeostasis and immune surveillance in synovial tissues, and its dysfunction is implicated in the progression of rheumatoid arthritis and osteoarthritis. Zbtb16, a transcription factor involved in cellular differentiation that is downregulated in mice with inflammatory arthritis and lymphatic defects, has been linked to skeletal development and immune regulation, but its direct role in lymphatic function remains unexplored. To investigate this, we generated Zbtb16^fl/fl^ mice for conditional-inducible loss-of-function experiments. Initial validation studies demonstrated that CMV-Cre x Zbtb16^fl/fl^ mice display the same skeletal phenotypes as previously reported Zbtb16^-/-^ global knockout mice. To achieve telocyte-specific deletion, Efhd1-CreERT2 x Zbtb16^fl/fl^ mice were treated with tamoxifen, and successful recombination was verified by PCR analysis of genomic DNA from popliteal lymphatic vessel tissue. Functional assessment using NIR-ICG imaging demonstrated that Zbtb16 deletion in telocytes partially impaired lymphatic clearance, suggesting a role in maintaining lymphatic function during homeostasis. These findings demonstrate the utility of Zbtb16^fl/fl^ mice for their intended purpose and support future studies on the role of Zbtb16 within the synovial lymphatic system.

## INTRODUCTION

The promyelocytic leukemia zinc finger (PLZF) protein, encoded by the *Zbtb16* gene, was originally identified in a patient with acute promyelocytic leukemia (APL) carrying the t(11;17)(q23;q21) chromosomal translocation that generates the PLZF-RARα and RARα-PLZF fusion proteins (*1, 2*). Both fusion products act as dominant-negative regulators of their native counterparts, contributing to leukemogenesis (*3*). Structurally, the Zbtb16 protein contains an N-terminal BTB domain and C-terminal Krüppel-type zinc fingers that enable dual transcriptional functions. Through the BTB domain, Zbtb16 recruits Polycomb-group proteins such as BMI1 and associated histone deacetylases (HDACs) to discrete nuclear foci (*4*), thereby enforcing transcriptional repression by modifying local chromatin conformation (*5*). Fusion of the Zbtb16 BTB domain to RARα in APL maintains myeloid cells in an undifferentiated state by aberrant recruitment of nuclear corepressors and HDACs (*6*), rendering Zbtb16-mediated repression sensitive to HDAC inhibition (*7*). In addition to repression, Zbtb16 can also activate transcription: phosphorylated Zbtb16 cooperates with promyelocytic leukemia protein (PML) and HDAC1 to bind interferon-stimulated gene (ISG) promoters and drive their transcription in response to IFN signaling (*8*).

Beyond its initial discovery in leukemia, Zbtb16 plays central roles in developmental biology. It regulates the balance between stem cell self-renewal and differentiation, including adult germline stem cell maintenance in spermatogenesis (*9*), myeloid progenitor maturation during early hematopoiesis (*10*), and proper axial skeleton and limb bud patterning (*11*). Zbtb16 also contributes to musculoskeletal development by binding developmental enhancers and promoting histone modifications such as H3K27 acetylation, thereby activating osteogenic gene programs required for human mesenchymal stem cell differentiation (*12*). Consistent with these regulatory functions, Zbtb16-deficient mice exhibit marked skeletal defects linked to dysregulated Hox gene expression (*11*). In pathological contexts, Zbtb16 has also been identified as a tumor suppressor and regulator of apoptosis and cell-cycle control, further indicating its importance in tissue maintenance and repair (*13*), further indicating its importance in tissue maintenance and repair.

Zbtb16 also operates as a context-dependent integrator of immune signaling. In interferon signaling pathways, its interaction with PML and HDAC1 is required for inducing a subset of interferon stimulatory genes (*8*). More broadly, Zbtb16’s chromatin-modifying activity positions it as a transcriptional hub capable of coordinating developmental, immune, and stress-response gene programs.

Lymphatic vessels are essential for tissue fluid balance and immune cell trafficking, particularly within synovial tissues (*14*). Impaired lymphatic drainage is a hallmark of inflammatory arthritis, where diminished lymphatic contractility and clearance amplify synovial inflammation and swelling (*15*). Although multiple stromal and immune cells contribute to the synovial lymphatic system (*16*), recent studies highlight telocytes (*2*) as a specialized interstitial population that maintains peri-lymphatic vessel structure and function. Yet, the transcriptional mechanisms governing telocyte-mediated lymphatic regulation remain unresolved. Our previous single-cell RNA-sequencing (scRNAseq) analysis of popliteal lymphatic vessel (PLV) tissue from WT and tumor necrosis factor-transgenic (TNF-tg) mice identified *Zbtb16* as one of the most significantly downregulated transcription factors in both telocytes and fibroblasts under chronic inflammatory conditions (*17*). Given its established roles in transcriptional control and tissue homeostasis, we aimed to test the hypothesis that loss of Zbtb16 in telocytes decreases their function and thereby impairs lymphatic drainage. To this end, here we describe the development and validation of Zbtb16^fl/fl^ mice, and preliminary lymphatic drainage studies with Zbtb16^fl/fl^ mice that were crossed with Efhd1-CreERT2 mice that allow tamoxifen-induced LoxP site recombination in telocytes.

## RESULTS

### Generation and Validation of Zbtb16 Conditional Knockout Mice

The Zbtb16 conditional knockout (Zbtb16^fl/fl^) strain was generated using CRISPR/Cas-mediated genome engineering as described in Figure 1 A. LoxP sites were inserted flanking exon 2 of *Zbtb16* in C57BL/6NTac mice to allow Cre-mediated excision. PCR genotyping confirmed the presence of LoxP-flanked alleles in founder mice, and sequencing analysis verified the precise insertion of LoxP sites without off-target mutations. These mice displayed no overt phenotypic abnormalities, confirming that insertion of LoxP sites did not disrupt gene function in the absence of Cre recombinase.

**Figure 1.**
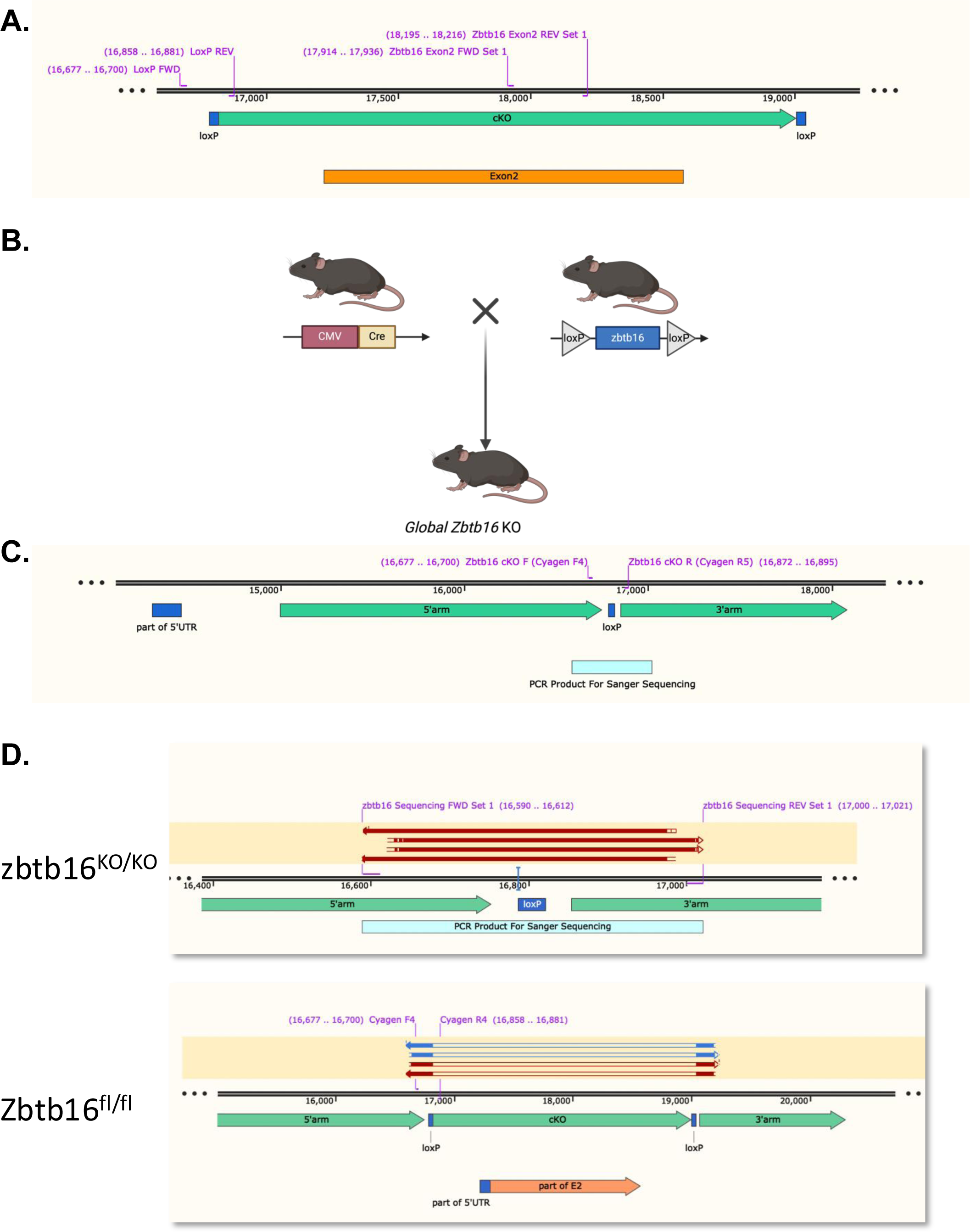
Generation and Validation of *Zbtb16* Conditional Knockout Mice. LoxP sites flanking exon 2 were inserted into the *Zbtb16* targeting construct to produce a conditional knockout allele in C57BL/6NTac mice using CRISPR/Cas-mediated genome engineering. To engineer the targeting vector, homologous arms and the cKO region were generated by PCR using BAC clone RP23-2I6 as template. Following sequencing validation, the targeting vector was co-injected with Cas9 and gRNA into fertilized eggs to produce Zbtb16^f/f^ mice. Genotyping of the pups was performed via PCR, followed by DNA sequencing to confirm the precise insertion loci (**A**). To validate this strain, Zbtb16^f/f^ mice were bred with CMV-Cre^+/−^ mice to generate global *Zbtb16* knockout mice (**B**) predicted to have the same skeletal phenotype as germline Zbtb16^-/-^ mice (*11*). LoxP recombination was designed to delete 62.8% of the *Zbtb16* coding region and induce a frameshift in the gene, which was confirmed by DNA sequencing (**C**). (**D**) Sanger sequencing was performed on genomic DNA from global Zbtb16^-/-^ mice to verify successful deletion. The sequencing results confirmed that both forward and reverse PCR products lacked the targeted cKO region, demonstrating effective Cre-mediated excision.

To confirm that the Zbtb16 ^fl/fl^ allele enables successful Cre-mediated recombination and gene deletion to produce the predicted null phenotype (*11*), we generated global Zbtb16 knockout (Zbtb16^-/-^) mice by crossing Zbtb16 ^fl/fl^ mice with CMV-Cre^+/−^ mice (Figure 1 B-C). Sanger sequencing of genomic DNA was performed on PCR products generated using primers targeting sequences flanking the cKO region in both the forward and reverse directions (Figure 1 D). Sequencing alignment confirmed the absence of the targeted exon 2 region in knockout mice, consistent with successful Cre-mediated recombination and excision of the floxed allele. The complete loss of the cKO region in both amplification directions provides molecular validation that the Zbtb16 ^fl/fl^ model functions as expected.

### CMV-Cre x Zbtb16^fl/fl^ mice Display the Zbtb16 Null Skeletal Phenotype

Given that conventional Zbtb16 knockout mice have a very specific skeletal defects (*11*), we tested to see if CMV-Cre x Zbtb16^fl/fl^ mice have the same phenotype. X-ray imaging and µCT confirmed that this strain develops the predicted tibia-fibula separation, indicating a loss of normal fusion (Figure 2A). Some CMV-Cre x Zbtb16^fl/fl^ mice also displayed a homeotic transformation of the tibia into a fibula-like structure, as well as the absence of the tibiale (Figure 2B), and homeotic transformation of the talus into a calcaneus-like structure (Figure 2 C-D). These abnormalities were absent in Zbtb16^fl/fl^ control littermates, verifying that Cre-mediated excision resulted in a functionally null allele, supporting the reliability of this strain for tissue-specific conditional-inducible Zbtb16 gene knockout studies.

**Figure 2.**
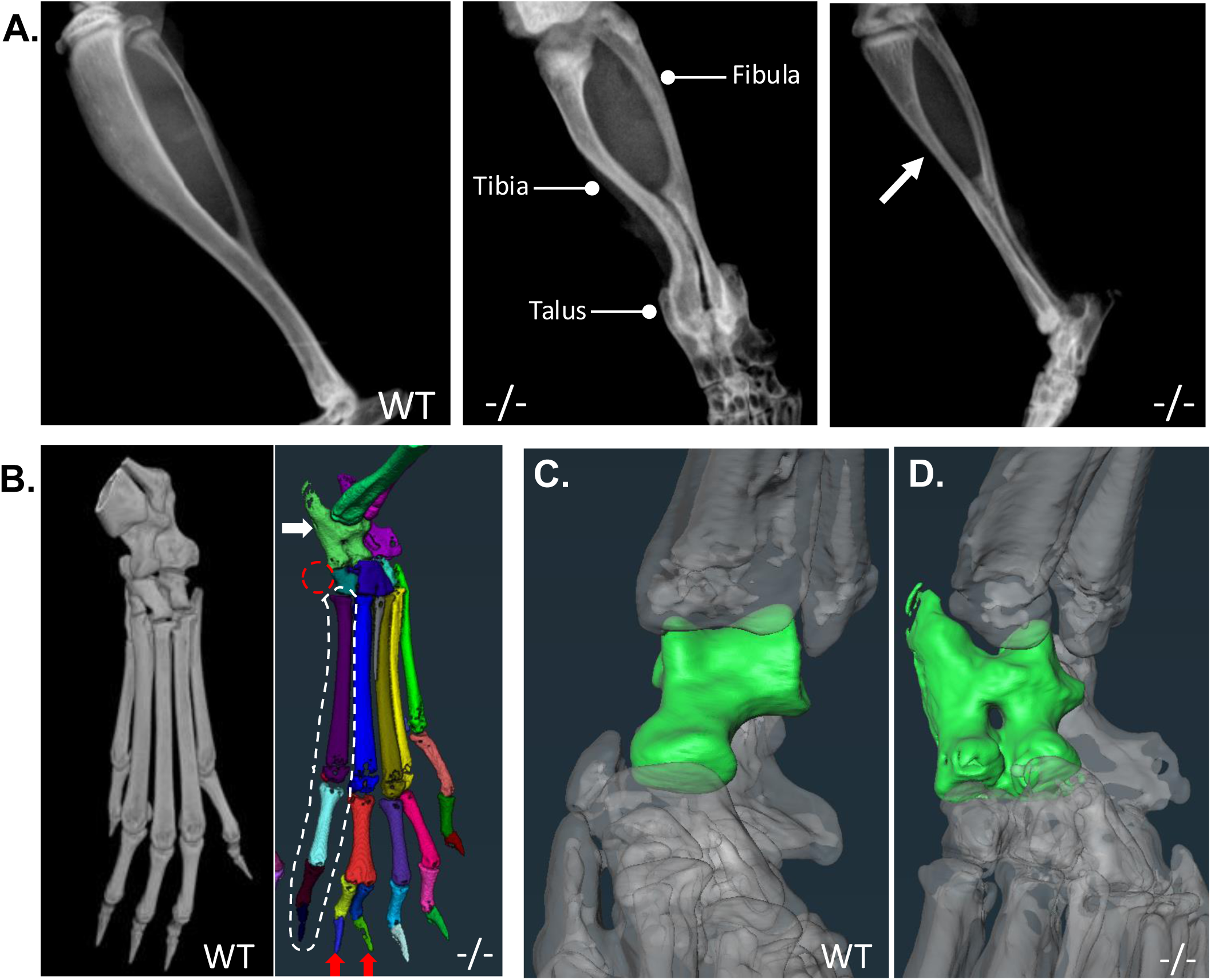
Validation of the *Zbtb16* Null Skeletal Phenotype in CMV-Cre x Zbtb16^f/f^ Mice. **(A)** X-rays of the lower limb confirm the known Zbtb16^-/-^ skeletal phenotype of CMV-Cre x Zbtb16^f/f^ (-/-) mice. Note the normal tibia and fibula successfully fused in WT Zbtb16^f/f^ mice (**left**). In contrast some -/- mice display separated/unfused tibia and fibula (**middle**), while in other mutant mice the tibia has undergone homeotic transformation into a fibula-like structure (**arrow in right**). (**B**) Micro-CT was performed on hindlimbs of WT and -/- mice and representative 3D renderings are presented to illustrate the homeotic transformation of the talus to calcaneus (white arrow), the absence of tibiale (red circle), the additional phalanx in digit I (white circle), and in some cases, the extra digit preaxially in digit II (red arrows) in the mutant mice. Comparison of talus micro-CT images from 7 weeks old WT (**C**) and -/- mice (**D**), highlighting the abnormal calcaneus-like talus in CMV-Cre x Zbtb16^f/f^ mice.

### Validation of Tamoxifen-Induced *Zbtb16* Gene Deletion in Peri-PLV Telocytes

As Efhd1-CreERT2 mice allow for gene-targeting in peri-PLV telocytes (*2*), we crossed the Zbtb16^fl/fl^ mice to this strain, and administered tamoxifen to induce Cre-mediated recombination (Figure 3A). Genomic DNA was extracted from PLV tissue, which contains a mixture of telocytes, adipocytes, and other stromal cells, and used it for PCR genotyping using primers designed to amplify both the floxed Exon 2 region (303 bp) present in all cells and the recombined conditional knockout (cKO) allele (219 bp), which is expected only in telocytes upon successful Cre-mediated recombination. The results showed that the Exon 2 band was detected in all samples, confirming the presence of *Zbtb16* genomic DNA across different cell populations. However, the cKO band was detected exclusively in PLV samples from Efhd1-CreERT2 x Zbtb16^fl/fl^ mice following tamoxifen treatment, but not in Cre-negative controls (Figure 3B). This indicates that Cre-mediated recombination and exon 2 deletion occurred specifically in telocytes. The absence of the cKO band in Cre-negative mice further confirms that recombination is dependent on the presence of CreERT2 expression in telocytes and tamoxifen induction.

**Figure 3.**
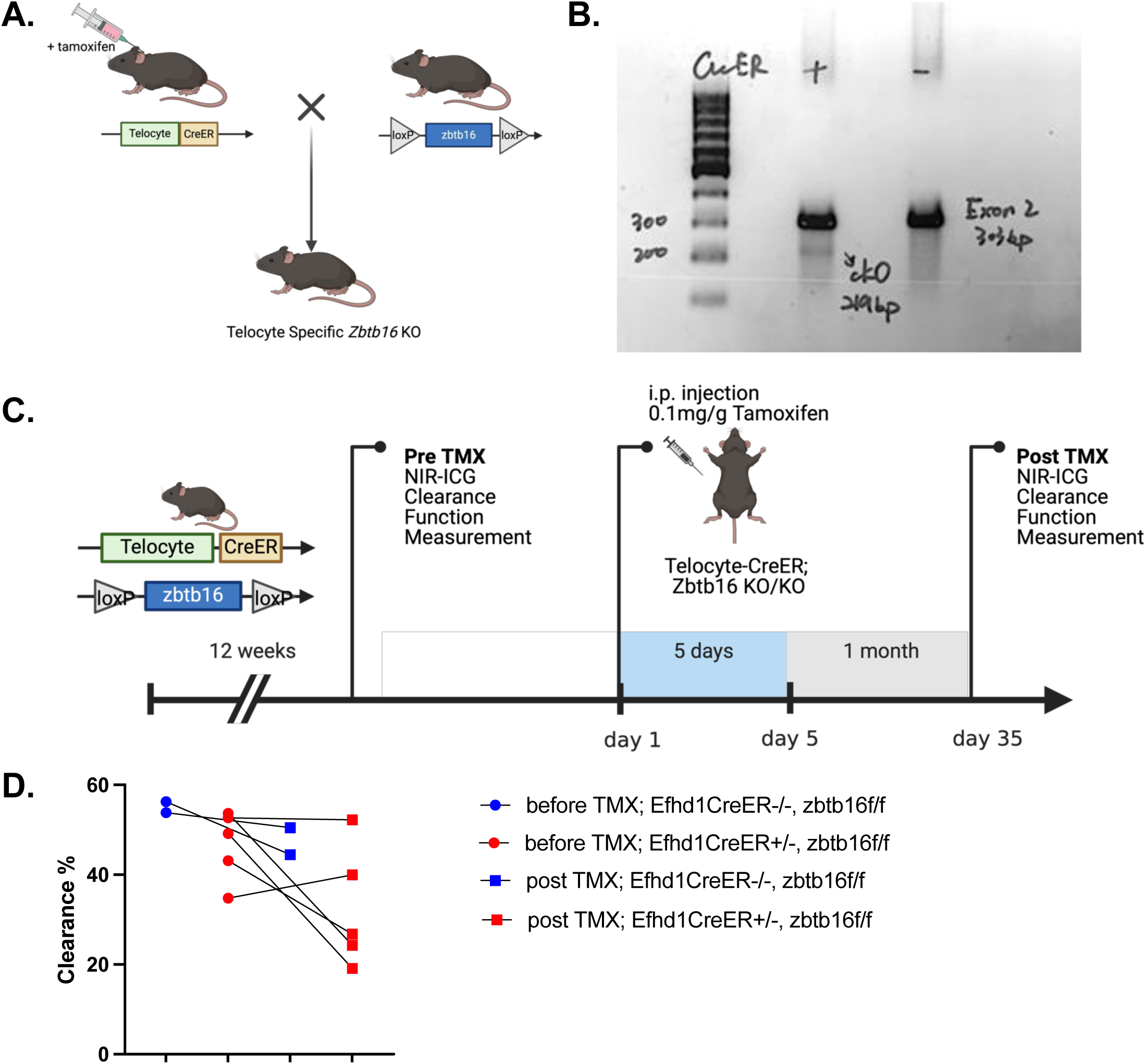
Telocyte-specific deletion of Zbtb16 partially exacerbates lymphatic dysfunction. (**A**) Schematic representation of the generation of telocyte-specific Zbtb16 knockout mice. Efhd1-CreERT2 mice were crossed with Zbtb16^fl/fl^ mice to obtain Efhd1-CreERT2 x Zbtb16^fl/fl^ offspring. (**B**) PCR analysis of PLV tissue confirmed successful Cre-mediated recombination in Efhd1-CreERT2 x Zbtb16^fl/fl^ mice, as indicated by the presence of the cKO band (219 bp) in Cre-positive samples. (**C**) Experimental timeline for inducing Zbtb16 deletion and assessment of lymphatic function via NIR-ICG imaging. (**D**) Quantification of lymphatic clearance function in WT (blue) and Efhd1-CreERT2 x Zbtb16^fl/fl^ (red) mice pre and post tamoxifen treatment, showing partial impairment in mutant mice at 1-month post-tamoxifen induction.

### Telocyte-Specific *Zbtb16* Knockout Decreases Lymphatic Clearance

To assess the functional consequences of telocyte-specific *Zbtb16* deletion, lymphatic clearance efficiency was evaluated using NIR-ICG imaging. Before tamoxifen introduction, no significant differences in lymphatic function were observed between control Zbtb16^fl/fl^ and Efhd1-CreERT2 x Zbtb16^fl/fl^ mice. However, one-month post-tamoxifen treatment, some cKO mice exhibited impaired clearance of the injected ICG dye from the hindlimb footpad, as indicated by increased residual fluorescence intensity compared to controls, suggesting that telocyte Zbtb16 plays a regulatory role in maintaining normal lymphatic vessel function.

## Discussion

Although there have been major advances in our understanding of lymphatic vessel biology (*18–21*), several critical knowledge gaps remain including the identity of cells that sense acute edema in tissue distal to collecting lymphatic vessels (cLV) to trigger enhanced lymphatic clearance by initiating pacemaking signals in lymphatic muscle cells (LMCs), and the mechanisms responsible for the loss of cLV contractility in lymphedema (*22, 23*) and arthritis (*15*). To fill this knowledge gap, we recently described the critical role of peri-PLV mast cells and telocytes in lymphatic clearance and joint homeostasis (*2, 24*). To expand on this and our finding that Zbtb16 is significantly downregulated in PLV tissue from TNF-tg mice will lymphatic defects and inflammatory-erosive arthritis (*17*), we generated a novel Zbtb16^fl/fl^ mouse line for cKO studies. Following validation of the genetics and phenotype of this line (Figures 1 and 2), we completed initial lymphatic drainage studies in Efhd1-CreERT2 x Zbtb16^fl/fl^ mice that suggest telocyte Zbtb16 contributes to the regulation of lymphatic vessel function (Figure 3).

One potential explanation for the relatively mild lymphatic dysfunction observed in Efhd1-CreERT2 x Zbtb16^fl/fl^ mice is that telocytes may have compensatory mechanisms under steady-state conditions. Transcription factors often exert their effects in response to environmental stimuli, and their impact may not be fully evident under basal conditions. Given that our previous work demonstrated that *in vivo* depletion of telocytes decreases lymphatic function(*2*), Zbtb16 may play a more significant role in telocyte-mediated lymphatic regulation during inflammatory or acute challenge conditions.

To better evaluate the functional impact of Zbtb16 deletion in telocytes, future studies could utilize an acute inflammatory arthritis model, such as zymosan-induced arthritis (ZIA) model (*2*). ZIA provides a controlled inflammatory environment that could reveal lymphatic defects that are not apparent under normal conditions. If Zbtb16 is required for telocyte adaptation during inflammation, we would expect Efhd1-CreERT2 x Zbtb16^fl/fl^ mice to exhibit exacerbated lymphatic dysfunction in response to ZIA, similar to preliminary data with Efhd1-CreERT2^+/−^ x DTA^f/-^ mice with ZIA(*2*).

Overall, our findings demonstrate that Zbtb16^fl/fl^ mice function as intended for cKO studies, and support a role for Zbtb16 in telocyte-mediated lymphatic function. However, further studies are needed to determine its precise mechanism of action. Investigating the effects of Zbtb16 deletion in inflammatory challenge models will be critical for understanding its contribution to lymphatic adaptation and its potential relevance in arthritis progression.

## Methods

### Mouse models

All animal research was conducted with approval by the University of Rochester Institutional Animal Care and Use Committee.

### Generation of Conditional Knockout Strain of Zbtb16

Zbtb16^fl/fl^ mice were generated using CRISPR/Cas-mediated genome engineering in C57BL/6NTac mice (Taconic Biosciences Inc.). LoxP sites were inserted flanking exon 2 of Zbtb16 to create a conditional knockout (cKO) allele. The targeting vector was designed with homologous arms and the cKO region, amplified using BAC clone RP23-2I6 as a template. Cas9, gRNA, and the targeting vector were co-injected into fertilized eggs to produce Zbtb16^fl/fl^ mice. Genotyping was performed using PCR, followed by DNA sequencing to confirm correct insertion. To validate the cKO strain, Zbtb16 ^fl/fl^ mice were crossed with CMV-Cre^+/−^ mice to generate global Zbtb16 knockout mice for phenotypic confirmation. For information of primers, see Table 1.

**Table 1.**
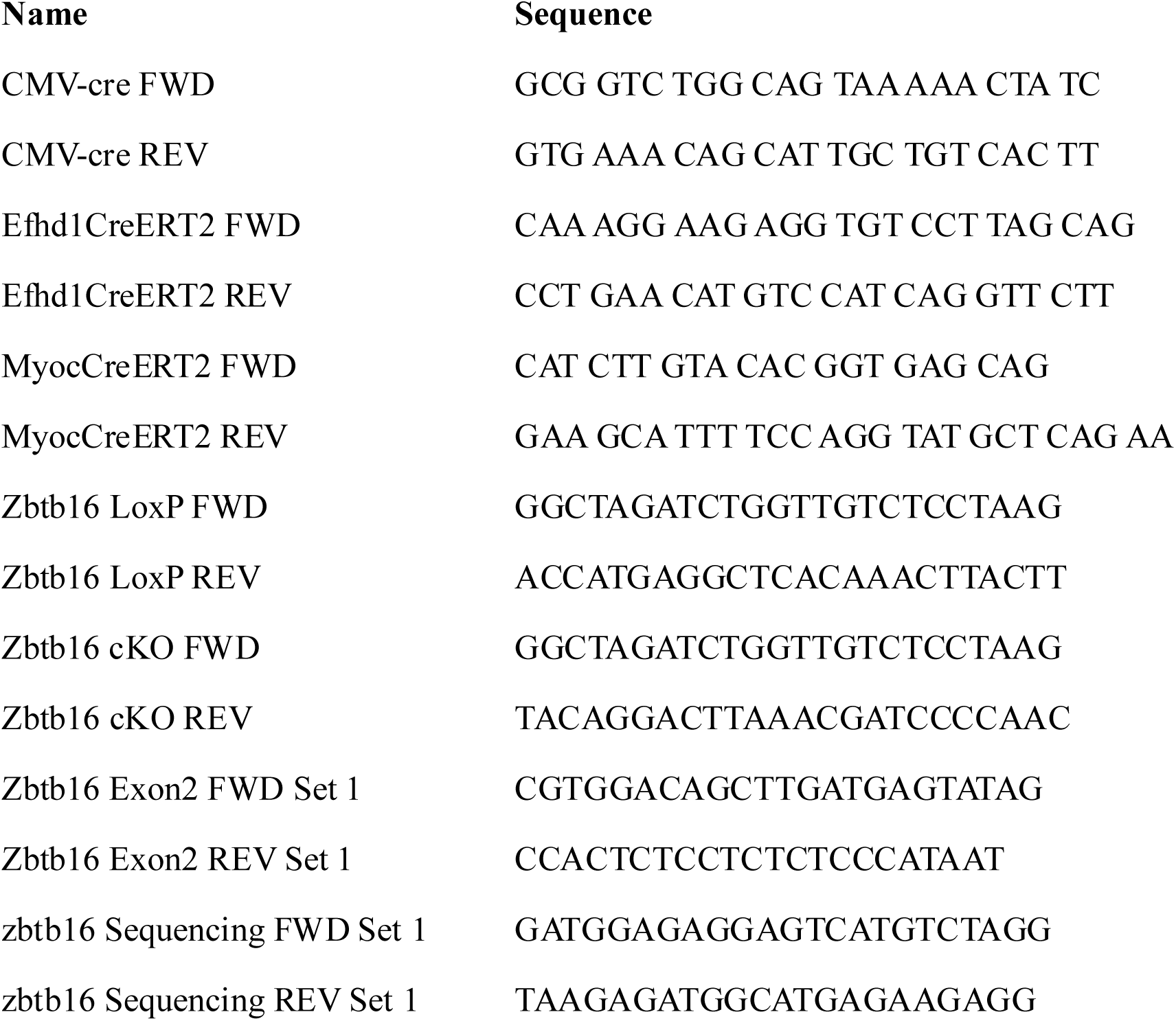
List of Primers.

### Generation of Telocyte-Specific Zbtb16 Knockout Mice

To generate telocyte-specific Zbtb16 knockout mice, Zbtb16 ^fl/fl^ mice were crossed with Efhd1-CreERT2 mice (*2*). Offspring carrying the Efhd1-CreER allele and homozygous floxed Zbtb16 (Efhd1-CreER x Zbtb16 ^fl/fl^) were treated with tamoxifen (100 mg/kg) via intraperitoneal injection for five consecutive days at 12 weeks of age to induce Cre-mediated recombination and Zbtb16 deletion in telocytes.

### Dual energy X-ray absorptiometry (DEXA)

Skeletal phenotyping of Zbtb16 KO and WT mice was performed using X-ray imaging (*25*) and µCT (*26*) as previously described. Mice were anesthetized, and X-ray images of the lower limbs were captured to assess skeletal abnormalities.

### Micro-CT analysis

For µCT analysis, hindlimbs were fixed in 10% neutral buffered formalin for 3 days, scanned at 10 μm resolution using a high-resolution µCT scanner, and reconstructed using Amira software to visualize bone morphology and homeotic transformations, as previously described(*26*), using identical imaging settings and parameters.

### NIR-ICG Lymphatic Clearance Assay Timeline and Data Collection

To assess lymphatic function, a fluorescent tracer-based lymphatic clearance assay was performed. Indocyanine green dye was injected into the footpad of hindlimb as previously described(*24*). NIR-ICG clearance function were measured pre- and 1-month post-tamoxifen treatment using an IVIS Spectrum imaging system. Clearance efficiency was quantified by measuring residual fluorescence intensity in the footpad. Tamoxifen treatment was administered at 12 weeks of age, followed by arthritis induction via intra-articular injection at day 1 post-tamoxifen. Mice were monitored at two key time points: day 0 pre-tamoxifen induction and day 35 post-arthritis induction.

## Acknowledgements

We would like to thank the faculty and staff at the Genomics Research Center and Histology, Biochemistry, Electron Microscopy Resource in the Center for Advanced Resource Technologies, and Molecular Imaging Core at the University of Rochester Medical Center. This work was supported by funding from the National Institutes of Health: R01AR056702 (EMS), and P30 AR069655 (EMS).

